# Particle Dynamics and Bioaerosol Viability of Aerosolized BCG Vaccine Using Jet and Vibrating Mesh Clinical Nebulizers

**DOI:** 10.1101/2021.04.26.441516

**Authors:** Rachel K. Redmann, Deepak Kaushal, Nadia Golden, Breeana Threeton, Stephanie Z. Killeen, Philip J. Kuehl, Chad J. Roy

## Abstract

**Background:** Bacillus Calmette–Guérin (BCG) is a vaccine used to protect against tuberculosis primarily in infants to stop early infection in areas of the world where the disease is endemic. Normally administered as a percutaneous injection, BCG is a live, significantly attenuated bacteria that is now being investigated for its potential within an inhalable vaccine formulation. This work investigates the feasibility and performance of four jet and ultrasonic nebulizers aerosolizing BCG and the resulting particle characteristics and residual viability of the bacteria post-aerosolization.

**Methods:** A jet nebulizer (Collison) outfitted either with a 3- or 6-jet head, was compared to two clinical nebulizers, the vibrating mesh Omron MicroAir and Aerogen Solo devices. Particle characteristics, including aerodynamic particle sizing, was performed on all devices within a common aerosol chamber configuration and comparable BCG innocula concentrations. Integrated aerosol samples were collected for each generator and assayed for bacterial viability using conventional microbiological technique.

**Results and Conclusions:** A batch lot of BCG (Danish) was grown to titer and used in all generator assessments. Aerosol particles within the respirable range were generated from all nebulizers at four different concentrations of BCG. The jet nebulizers produced a uniformly smaller particle size than the ultrasonic devices, although particle concentrations by mass were similar across all devices tested with the exception of the Aerogen Solo, which resulted in a very low concentration of BCG aerosols. The resulting measured viable BCG aerosol concentration fraction produced by each device approximated one another; however, a measurable decrease of efficiency and overall viability reduction in the jet nebulizer was observed in higher BCG inoculum starting concentrations, whereas the vibrating mesh nebulizer returned a remarkably stable viable aerosol fraction irrespective of inoculum concentration.

## Introduction

Tuberculosis (Tb) is a worldwide endemic pulmonary disease that infects 1/3 of mankind and is the causative disease agent of over two million deaths yearly (1, 2). Tubercular disease in humans can be attributed to respiratory exposure to and ensuing infection from *Mycobacterium tuberculosis*, an environmentally hardy pathogenic mycobacteria that has caused human disease for several thousand years (3). A prophylactic vaccine that protects people of all ages against infection has been a goal of medical science for several decades. Many types of Tb vaccines have since been developed (1, 4), which include live attenuated, whole cell killed, virally-vectored, and subunit variants. One of the earliest attempts at vaccine development yielded a live, highly attenuated multi-passage variant of *Mycobacterium bovis* referred to as Bacillus Calmette–Guérin (BCG), accrediting the creation to its French originators. BCG remains the only licensed vaccine for prevention of tubercular infection and is still in use today (5–7). BCG has been widely utilized for several decades primarily in pediatric populations in countries where Tb is considered endemic. BCG is currently administered as a subcutaneous injection and has proven to be effective in the prevention of childhood pulmonary Tb when administered to infants. The vaccine is considered to result in minimal efficacy, however, when administered to adults and has not proven to be effective in the generation of an immune response that is consistent with protection against pulmonary Tb (5).

Several years ago, it was proposed that the apparent lack of stimulating the appropriate immune response in the adult that would provide protection against tubercular disease may be associated with the route of vaccination rather than the biological components and composition of the vaccine being used (8–10). The strategy of delivering BCG as an aerosol or mucosally to stimulate preferential immune response has been performed in animals and clinically for a number of years (9, 11–13). *M. tuberculosis* is an exceedingly complex microorganism that has adapted to the rare ontological niche of being a successful human pathogen over thousands of years. It stands to reason that a successful vaccine to Tb would by necessity contain the majority of the biological complexities associated with the disease agent, and maintain the structural and biological capacity for replication which are many of the characteristics of BCG (7, 14, 15). In addition, Tb is an obligate respiratory pathogen - the primary (and required) route of infection for *M. tuberculosis* is through inhalation of infectious aerosols sourced from an infected host and/or fomite re-aerosolization (16–21). An aerosolized form of BCG, therefore, would maintain the requirement for a ‘whole’ vaccine product thought to be one of the necessities for adult protection, while delivering to the respiratory system, eliciting an immune response both humoral and locally in lung mucosa, which would provide dual protection against disease. Immune response of mucosally-delivered BCG has been investigated over the past several years (9, 12, 22) with some success when used as a comparator to conventional immunization strategies.

Aerosol generation of a biologically active, replication-competent microorganism such as BCG necessitates characterization of the device provisioning the aerosol in order to define physical composition and immunizing ‘dose’. Investigation of the physical and biological characteristics of the aerosols produced and the resulting effects of the corresponding mycobacterial payload within the particles is essential to this process. Accordingly, in this work we assess the effects of aerosolization upon BCG when using various aerosol generators. Initially, particle size dynamics of BCG was assessed at four discrete starting concentrations, hereafter referenced to notationally as *C_s_* (at 10^5^, 10^6^,10^7^,10^8^ CFU/ml) when using either jet or vibrating mesh nebulizers. Particle size characteristics were assessed using a singular method, aerodynamic time of flight measurements by an aerodynamic particle sizer (Thermo Systems Inc (TSI), St. Paul, MN). Two primary parameters including the mass median aerodynamic diameter (MMAD) and count median aerodynamic diameter (CMAD) and corresponding aerosol concentration by mass and number were determined from measurements performed using the APS. These parameters were measured using four discrete starting bacterial concentrations of BCG; all BCG aerosol events were performed using the same growth lot of BCG bacteria. Thereafter, in discrete set of aerosol experiments, integrated sampling for the purposes of quantifying aerosolized culturable bacteria was performed using liquid impingement. The results of the impingement provided calculated aerosol concentrations of culturable BCG from each nebulizer for each bacterial starting concentration used in the study. Collectively, the physical characterization and particle sizing profile paired with the culturable mycobacterial aerosols provides a physical and biological basis from which to determine dose as a function of inhalation of generated aerosol.

## Materials and Methods

### Nebulizers

The jet nebulizers that were used in this characterization are either 3- or 6-jet Collison nebulizers (CH Technologies, Westwood, NJ). This style nonclinical nebulizer provides input feed through Bernoulli effect capillary uptake from a liquid reservoir and entrains the liquid into an provided airstream that function at critical flow (2 lpm/jet) under pressure; a minimal 18 psig is required to run the complementary airstreams to achieve optimal particle size (23). The 3-jet Collison nebulizer operates at 6 lpm; the 6-jet at 12 lpm. The mesh nebulizers used for comparison to the jet nebulizers include the Omron Microair and Aerogen solo. Both versions are employed in clinical settings for the delivery of aerosolized medication. The generator platform, rather than using jet airflow, utilizes a mesh (palladium) perforated with conical holes that act as a micropump when vibrated (24). Neither vibrating mesh generator uses intrinsic air flow as a means to deliver aerosol, rather the action of the vibrating mesh and respiratory inspiration works to deliver aerosols.

### Measurement with Aerodynamic Particle Sizer

Particle characteristics were determined using an aerodynamic particle sizer (APS Model 3321, TSI Inc., St. Paul, MN). The APS measures the aerodynamic size of particles from 0.5 - 20 microns and uses time-of-flight analysis based upon velocity and relative density of interrogated particle stream to determine particle behavior while airborne. Aerosol is drawn into the APS at a total flow of 5 l/min; 20% of the total flow is dedicated to inlet into the analyzer; 80% is sheath flow. The APS spectrometer uses a double-crest dual laser system and nozzle configuration which reduces the advent of false (e.g., doublet) background counts. Analysis of data from the APS was collected and device software (Aerosol Instrument Manager Version 5.3, TSI Inc., St Paul, MN) was used for initial review of data. Statistical analysis and graphing was performed using GraphPad (Prism V.7, GraphPad, La Jolla, California). The APS device operated on a continual basis once aerosol generation was initiated, and logged data for the duration of each discrete aerosol event.

### Experimental configuration

All BCG aerosols took place inside a 16-liter polycarbonate chamber outfitted with dilution and exhaust tubing and a sampling orifice. The chamber was connected to an automated system (Biaera Technologies, Hagerstown, MD) which controlled dilution, exhaust, sampling, and generator air flows when applicable, and also recorded temperature, relative humidity, and pressure readings. The automated system maintained equal rates of total air flow in and out of the chamber in order to retain equilibrium. **Figure 1** illustrates the experimental configuration of the chamber utilizing one of five possible nebulizers/ nebulizer orientations and one of two sampling strategies implemented. The jet nebulizer (A) used was either the 3-jet or 6-jet Collison, which require an outside air supply source, provided by the automated system in this instance. The handheld mesh nebulizers, however, are electronically operated and do not require outside air supply. They were placed inside the chamber and clamped to a shelf to achieve the proper height for proximity to the sampler tubing within the chamber. The mesh nebulizers used in this study were the (B) Omron Microair held horizontally, the (C) Omron Microair tilted 30° from horizontal, and the (D) Aerogen Solo. Samples from the chamber were collected using either the (E) APS for particle characterization or the (F) All Glass Impinger (AGI) sampler for bacterial viability. Total air flow in and out of the system was kept at 16 l/min, with adjustments to the dilution and exhaust flows as needed for differing generator and sampling requirements.

**Figure 1.**
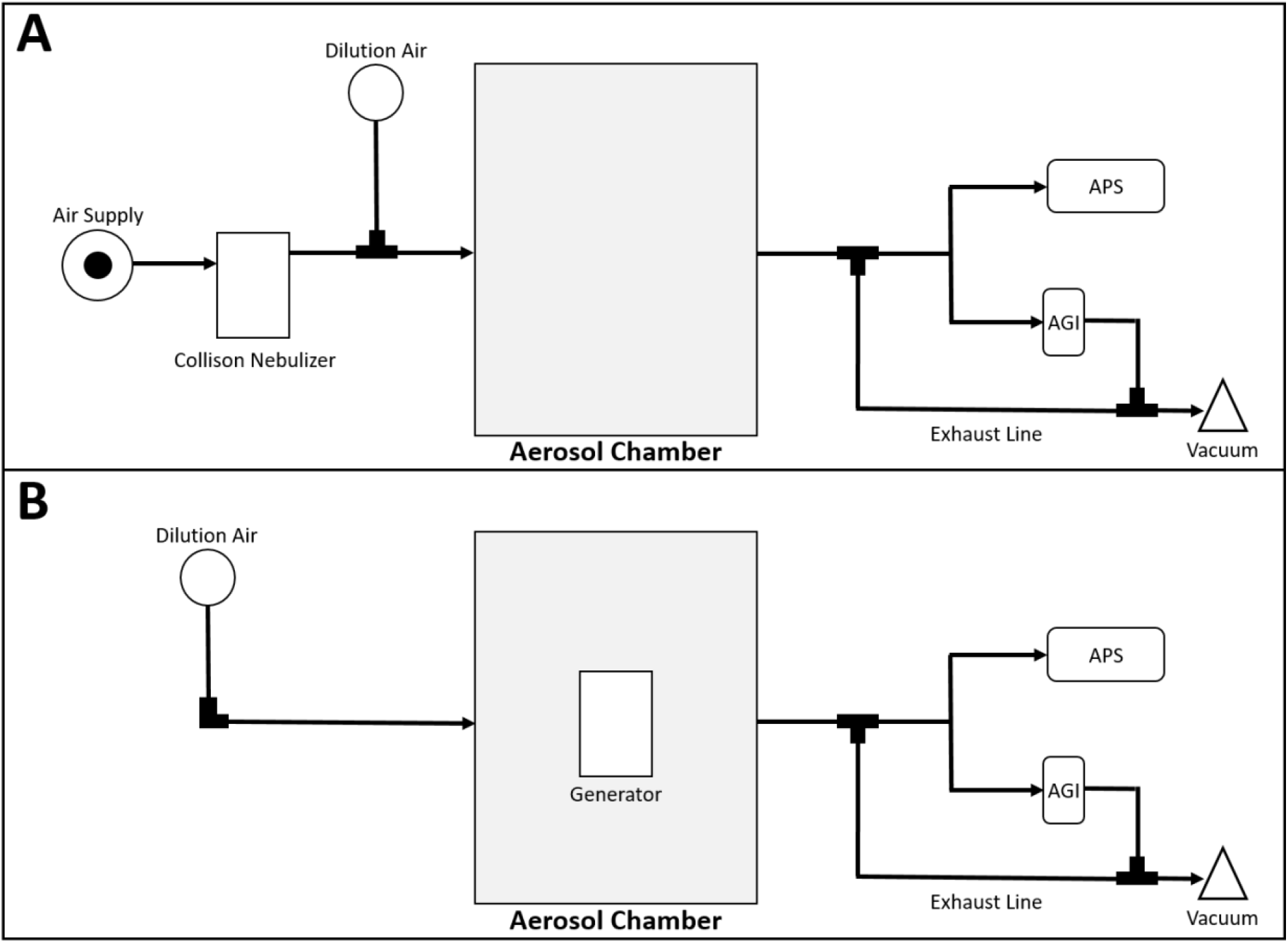
Experimental setup and corresponding sampling strategy used for the evaluation of nebulizers. BCG was aerosolized using a 3-jet or 6-jet Collison nebulizer (panel A), or using the Omron or Aerogen vibrating mesh aerosol generators in a separate configuration (panel B). Aerosol samples were collected using the APS to determine particle size dynamics and the AGI-4 Sampler for the purposes of bioaerosol viability in both configurations.

### Experimental Procedure

Individual aliquots of a liquid volume of BCG (5 ml) was prepared for each device and corresponding dilution within assessment for each nebulizer device. Upon discrete performance for each device, a liquid aliquot was directly expressed into the precious fluid reservoir for the Collison nebulizers, the medication port for the Omron MicroAir device, and the reservoir basin for the Aerogen solo. Each device was then actuated, and allowed to continuously run for analysis. The configuration supporting device evaluation was harmonized between devices, and shared similar design and internal volume (**Figure 1**). Flow rates for the configured system varied according to the device being used to accommodate for the relatively high flow rates generated by the Collison nebulizers (6 and 12 LPM for the 3-jet and 6-jet versions, respectively) compared to the devices with no intrinsic air flow (Omron MicroAir and Aerogen Solo) which relies upon patient inspiratory flow to facilitate aerosol delivery. Therefore, input flow for the Omron and Aerogen devices in this configuration was augmented with an external pump that provided equivalent input flow into the chamber at rate approximating the 3-jet Collison (6 LPM). Two aerosol sampling devices with differing flows were used in discrete aerosol generation events. For the experiments involving PSD, the aerodynamic particle sizer (TSI Model 3321) was used which houses an internal exhaust flow of 5 LPM. Residual exhaust flow was provided via an external pump at 2 LPM. BCG aerosols were also collected in discrete aerosol generation events for the purposes of biological viability determination of the BCG aerosols. The AGI sampler requires 6 LPM exhaust flow for operation, therefore the residual exhaust flow from the chamber was adjusted according to each device requirement and the necessity to maintain neutral pressure (0” H20) which was actively monitored throughout every aerosol generation event. The dynamic flows as described through the evaluation chamber were operated continuously for every evaluation for each device. Temperature and humidity was monitored during all evaluations. The prevailing temperature was 21.6±2.6° C and relative humidity 51.5±8.3% across all evaluations.

### Propagation and quantifying BCG

The vaccine BCG (Danish) was commercially acquired through ATCC (Manassas, VA). The stock vial, which was held at −80° C, was thawed at RT and 100 μl added to 10 ml of Middlebrook 7H9 media (Fisher Scientific, Hanover Park, IL), warmed to 37° C, then agitated on an incubated shaker for approximately 3 days. Subcultures were grown at a 1:10 ratio of subculture aliquot to Middlebrook 7H9 media (Fisher Scientific, Hanover Park, IL) until an OD of approximately 0.5 was attained. The initial subculture was derived from a stock culture, and all subsequent subcultures thereafter were propagated from a previous subculture. The stock culture and all subcultures were held at 4° C until aerosolization. Bacterial concentration was confirmed by plating 100 μL of the final subculture used for aerosolization, on prepared 7H11 agar (Fisher Scientific, Florence, KY) media plates. The stock culture was held at 4° C until subculture for the purposes of experimental use.

## Results and discussion

### PSD of BCG aerosols

The count median diameter (CMAD), mass median aerodynamic diameter (MMAD), and geometric standard deviation (GSD) for each measurement, shown in **Table 1**, represents the overall mean and corresponding standard deviations across all starting concentrations of BCG (10^5^-10^8^ CFU/ml) performed with each aerosol generator. Close examination of individual starting concentrations indicated little variation in particle size characteristics, and statistical comparison resulted in no significant differences (p>0.05) between BCG starting concentrations in particle characteristics when using the same aerosol generator.

**Table 1.** Particle sizing dynamics, mass median diameter, count median diameter, and corresponding geometric standard deviations for each nebulizer evaluated.

| *generator* | *MMAD ($\mu\text{m}$)* | *GSD ($\mu\text{m}$)* | *CMAD ($\mu\text{m}$)* | *GSD ($\mu\text{m}$)* |
| --- | --- | --- | --- | --- |
| 3JC | 2.69±0.03 | 1.71±0.01 | 1.20±0.02 | 1.48±0.00 |
| 6JC | 3.08±0.04 | 1.63±0.01 | 1.36±0.03 | 1.54±0.00 |
| OM | 2.76±0.06 | 1.59±0.01 | 1.41±0.07 | 1.50±0.00 |
| OM.t | 3.66±0.19 | 1.47±0.02 | 1.98±0.12 | 1.59±0.01 |
| AeS | 3.04±0.02 | 1.67±0.00 | 1.13±0.01 | 1.61±0.00 |

The CMAD across all aerosol generators was remarkably similar, and ranged from 1.1 to 1.6 μm and was not affected by starting concentration used in the aerosol generation events as evidenced by low heterodispersity GSD (range:1.48-1.59). Similarly, the MMAD for all generators were collectively below 4 μm (range:2.69-3.66) with little variation in the corresponding GSD (range:1.47-1.71), indicating minimal effect on the density solute of the aerosols generated when using higher concentrations of BCG. The majority of the particles represented in the corresponding distributions were below 5.8 μm, and are considered the fine particle fraction (FPF) of aerosols when collectively describing the characteristics of the distribution. Accordingly, the percentage of FPF represented as a part of the whole distribution was >90% for all generators evaluated using BCG.

There were differences in the number and mass of particles, measured as an airborne concentration, from each generator evaluated. The total number of particles, expressed as particles/cm^3^ of aerosol, and as a measure of mass generated, expressed as mg/m^3^, is shown in **Table 2**. Theoretical ‘dose’ of BCG, calculated based upon a series of inhalation presumptions, only considering viable fraction of BCG post-aerosolization, is shown in **Table 2**. Doses are shown stratified by the initial BCG concentration (in CFU/ml) and according to the nebulizer under evaluation. Three of the four nebulizers (3JC, 6JC, and OM) produced remarkably similar number and mass of particles generated, with the OM.t (Omron in a 30° orientation) producing noticeably more particles by mass than any other generator tested. The Aerogen Solo (AeS) produced the lowest number of particles by number and mass, returning a logarithmically lower (~0.327 mg/m^3^) mass concentration. Accordingly, the results of the theoretical calculated dose shown in **Table 2** is apropos as demonstration that only a small portion (<1% in many cases) of the post-aerosol culturable BCG is available for inhalation when considering the initial BCG (CFU/ml) concentration used in each nebulizer.

**Table 2.**
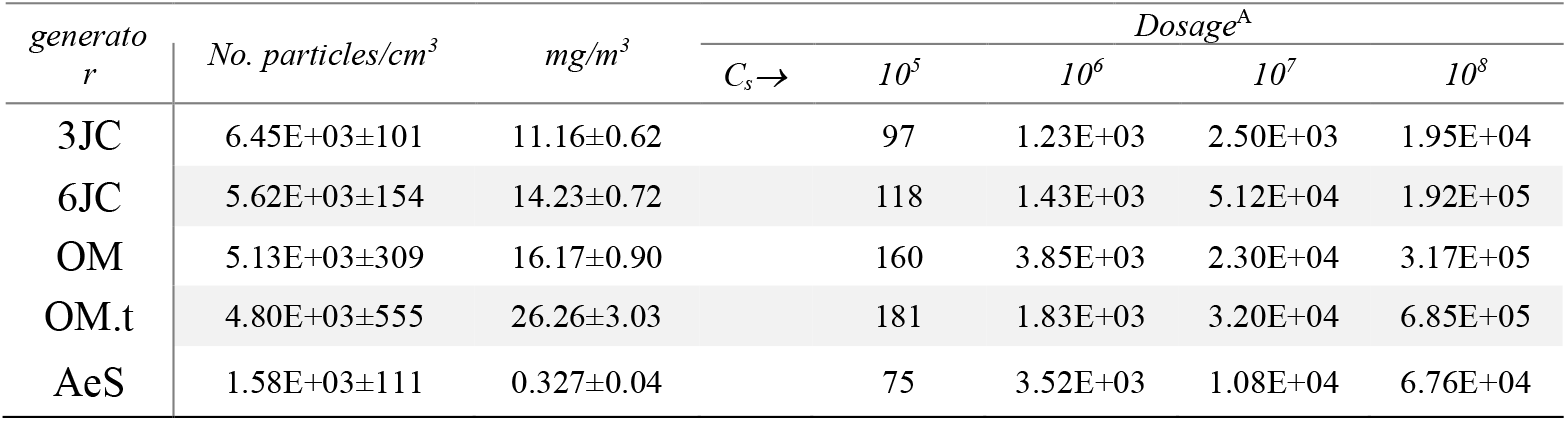
Aerosol concentration as a function of particles and mass and resulting dosage of viable BCG based upon predicted inhalation constants

Comparison of nebulizer performance, when assessed purely as an efficiency of total aerosol particles generated by a nebulizer, can be informative to overall contribution of the ‘viable fraction’ as a percentage of total number of particles generated. **Figure 2** details the percentage of BCG aerosol particles as a function of prevailing *C_s_* in use and total particles generated by each nebulizer under evaluation. All nebulizers were remarkably uniform in total number of particles generated (~5E+07 particles/liter of aerosol) with exception of the Aerogen Solo (~1E+07 particles/liter of aerosol) across all *C_s_* performed, stratified logarithmically. The relative percentage contribution of the viable BCG as a component of the total particles generated, which was calculated *post hoc* to analysis and functionally as *C_a_*, demonstrates significant differences between nebulizers assessed. For example, at a *C_s_* of 1E+07 CFU/ml, the relative percentage contribution of viable BCG as a component of total particles generated for the 3-jet Collison nebulizer was ~1E-04% compared to the Omron MicroAir, which showed the ~2E-02%, showing a 2-log difference in viability contribution at a *C_s_* of 1E+07. Differentials in viability and the overall performance of the relative efficiency of nebulizers can be summarized as Om.t>OM>AeS>6JC>3JC at the highest *C_s_* (1E+08).

**Figure 2.**
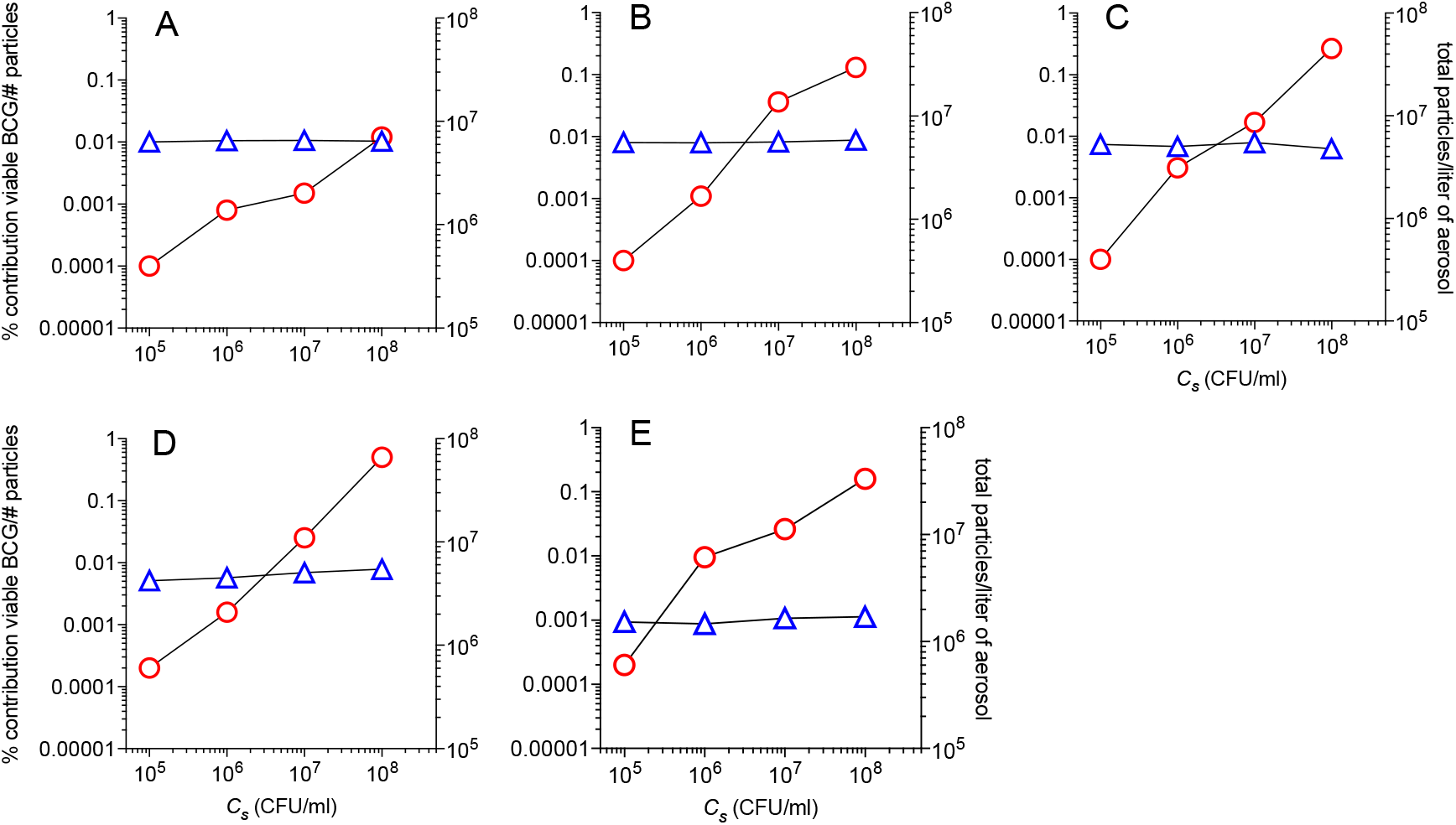
Comparison of nebulizer performance as a function of total particles per liter of aerosol generated, blue triangles (right ordinate axis), percentage contribution of viable BCG as a function of total aerosol particles generated, red circles (left ordinate axis) based upon BCG starting concentration (*C_s_*) used for each aerosol generation event (abscissa axis). Each panel represents A. 3-jet Collison, B. 6-jet Collison, C. Omron MicroAir, vertical orientation, D. Omron MicroAir, 30° vertical orientation, and E. Aerogen solo.

### Spray factor of BCG

The spray factor (*F_s_*) is a unitless ratio calculated by the prevailing inoculum loaded into the generator (*C_s_*) to the aerosol concentration determined to have been generated (*C_a_*), both of which are expressed as CFU per liter(23, 25–27). Calculation of *F_s_* is a useful quotient used to understand the dilution and relative effect of aerosol generator action upon viability of the particular biological agent under study. The results of the *F_s_* for BCG in each nebulizer evaluated, and stratified according to prevailing logarithmic *C_s_*, is shown in **Figure 3**. There are clear differences in *F_s_* between nebulizers under evaluation. The experimental determinations resulting from 3JC nebulizer demonstrated that initial *F_s_* at 1E+05 *C_s_* (3.1E-07) significantly worsened by nearly one log as BCG *C_s_* logarithmically increased to 1E+08 (1.3E-08). A similar trend was observed in the 6JC evaluation, with a 0.5log worsening of the *F_s_* between the lowest (1E+05) and highest (1E+08) *C_s_* performed. In contrast, the *F_s_* for OM, OM.t, and AeS is relatively stable as a function of BCG *C_s_* used in each discrete evaluation.

**Figure 3.**
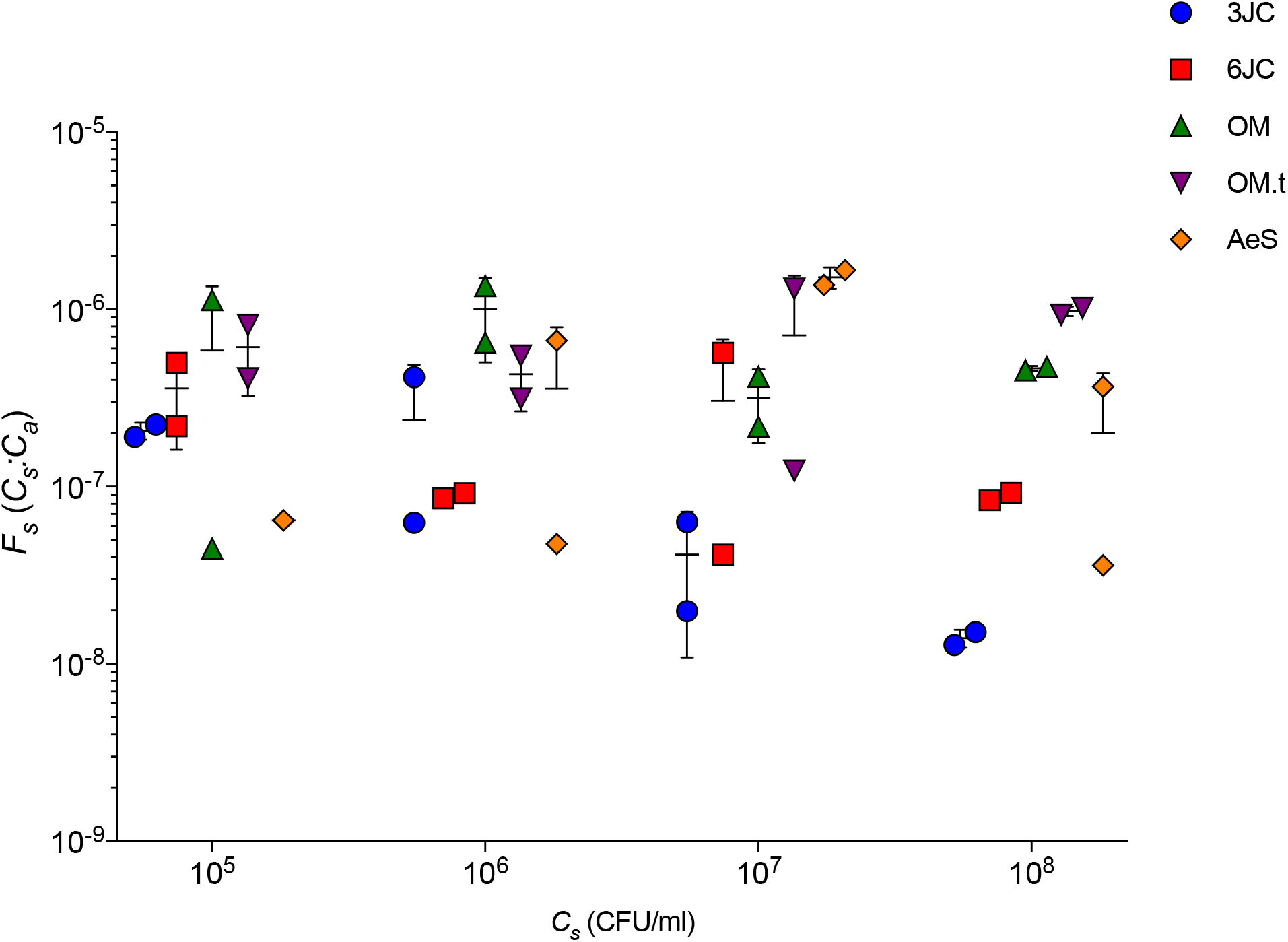
Spray factors (*F_s_*) for BCG. Corresponding lines show standard deviation and median of discrete experiments by aerosol generator and dilution of BCG inoculum used.

#### Effect of aerosolization on BCG

The mechanism by which each style nebulizer generates the aerosol affects the resulting viability of the BCG. The Collison nebulizer employs a multitude (either 3 or 6 jets) two-fluid nozzle jets under pressure that produces a high-velocity stream impacting the wall of the reservoir containing the BCG inoculum. The satellite aerosols from this action are swept up into the subsequent flow out of reservoir and outlet with a total flow of either ~6 (3JC) or ~12 (6JC) LPM. The mechanical shear developed during this process of nebulization undoubtedly imparts stress onto the mycobacterial innocula, and may further degrade the capacity of future culturability. The ultrasonic nebulizer (OM), in contrast, utilizes a piezoelectric actuated mesh in order to generate an aerosol. The Aerogen Solo (AeS) uses similar technology, with the exception of utilizing a palladium mesh for the purposes of aerosol generation. The surface tension of the liquid to be nebulized for both the OM and AeS pushes onto the metallic substrate via gravity flow to initiate generation, and there is no intrinsic air flow to carry the aerosol from the generator and rather relies upon flow from inhalation velocity of a patient or other corollary flow. Mechanical shear is minimized in both latter nebulizer designs. The distinction between the Collison (C) and OM/AeS nebulizers as it relates to the worsening *F_s_* as BCG *C_s_* increases (**Figure 3**) may be a result of the coarse treatment of the contents of the innocula in the former and relatively gentle single-pass generation method used by the OM and AeS nebulizers.

## Acknowledgments

Special thanks to the Aerosol and Mucosal Vaccination Community standing membership of the Bill and Melinda Gates Foundation (BMGF)-based Collaboration for Tuberculosis Vaccine Discovery (CTVD) for outstanding discussions and critical analysis of the subject matter of this work. The authors also acknowledge the Infectious Disease Aerobiology Core located at the Tulane National Primate Research Center, Tulane University, for logistical and scientific coordination and use of its aerobiology resources essential to this work.

## Funding Acknowledgement

This work was funded by the BMGF Grant No. OPP1126491 to CJR, the Aeras Foundation Innovation Fund/MISC041 to CJR and PK, and supported in part by Grant No. OD011104 to CJR from the Office of Research Infrastructure Programs (ORIP), Office of the Director, National Institutes of Health.

## Author Disclosure Statement

No competing financial interests exist.

## Notes

### Competing Interest Statement

The authors have declared no competing interest.

## REFERENCES

1. Cayabyab MJ, Macovei L, Campos-Neto A. Current and novel approaches to vaccine development against tuberculosis. Front Cell Infect Microbiol. 2012;2:154.

2. Raviglione MC. The new Stop TB Strategy and the Global Plan to Stop TB, 2006-2015. Bull World Health Organ. 2007;85(5):327.

3. Lienhardt C, Bennett S, Del Prete G, Bah-Sow O, Newport M, Gustafson P, et al. Investigation of environmental and host-related risk factors for tuberculosis in Africa. I. Methodological aspects of a combined design. Am J Epidemiol. 2002;155(11):1066–73.

4. Kamath AT, Fruth U, Brennan MJ, Dobbelaer R, Hubrechts P, Ho MM, et al. New live mycobacterial vaccines: the Geneva consensus on essential steps towards clinical development. Vaccine. 2005;23(29):3753–61.

5. Lagrange PH. [Successes and failures of BCG vaccination. Myths and realities. II]. Arch Fr Pediatr. 1982;39(4):271–4.

6. Roy A, Eisenhut M, Harris RJ, Rodrigues LC, Sridhar S, Habermann S, et al. Effect of BCG vaccination against Mycobacterium tuberculosis infection in children: systematic review and meta-analysis. BMJ. 2014;349:g4643.

7. Dockrell HM, Butkeviciute E. Can what have we learnt about BCG vaccination in the last 20 years help us to design a better tuberculosis vaccine? Vaccine. 2021.

8. Nagpal PS, Kesarwani A, Sahu P, Upadhyay P. Aerosol immunization by alginate coated mycobacterium (BCG/MIP) particles provide enhanced immune response and protective efficacy than aerosol of plain mycobacterium against M.tb. H37Rv infection in mice. BMC Infect Dis. 2019;19(1):568.

9. Manjaly Thomas ZR, McShane H. Aerosol immunisation for TB: matching route of vaccination to route of infection. Trans R Soc Trop Med Hyg. 2015;109(3):175–81.

10. Kaushal D, Foreman TW, Gautam US, Alvarez X, Adekambi T, Rangel-Moreno J, et al. Mucosal vaccination with attenuated Mycobacterium tuberculosis induces strong central memory responses and protects against tuberculosis. Nat Commun. 2015;6:8533.

11. White AD, Sarfas C, Sibley LS, Gullick J, Clark S, Rayner E, et al. Protective Efficacy of Inhaled BCG Vaccination Against Ultra-Low Dose Aerosol M. tuberculosis Challenge in Rhesus Macaques. Pharmaceutics. 2020;12(5).

12. Manjaly Thomas ZR, Satti I, Marshall JL, Harris SA, Lopez Ramon R, Hamidi A, et al. Alternate aerosol and systemic immunisation with a recombinant viral vector for tuberculosis, MVA85A: A phase I randomised controlled trial. PLoS Med. 2019;16(4):e1002790.

13. Maclouf AC, Fichez LF. [BCG aerosol vaccination]. Rev Tuberc. 1952;16(10-11):1016–8.

14. Palendira U, Spratt JM, Britton WJ, Triccas JA. Expanding the antigenic repertoire of BCG improves protective efficacy against aerosol Mycobacterium tuberculosis infection. Vaccine. 2005;23(14):1680–5.

15. Prentice S, Dockrell HM. Antituberculosis BCG vaccination: more reasons for varying innate and adaptive immune responses. J Clin Invest. 2020;130(10):5121–3.

16. Hella J, Morrow C, Mhimbira F, Ginsberg S, Chitnis N, Gagneux S, et al. Tuberculosis transmission in public locations in Tanzania: A novel approach to studying airborne disease transmission. J Infect. 2017;75(3):191–7.

17. Wood R, Morrow C, Barry CE, 3rd, Bryden WA, Call CJ, Hickey AJ, et al. Real-Time Investigation of Tuberculosis Transmission: Developing the Respiratory Aerosol Sampling Chamber (RASC). PLoS One. 2016;11(1):e0146658.

18. Jones-Lopez EC, White LF, Kirenga B, Mumbowa F, Ssebidandi M, Moine S, et al. Cough Aerosol Cultures of Mycobacterium tuberculosis: Insights on TST / IGRA Discordance and Transmission Dynamics. PLoS One. 2015;10(9):e0138358.

19. Shenoi SV, Escombe AR, Friedland G. Transmission of drug-susceptible and drug-resistant tuberculosis and the critical importance of airborne infection control in the era of HIV infection and highly active antiretroviral therapy rollouts. Clin Infect Dis. 2010;50 Suppl 3:S231–7.

20. Fennelly KP. Variability of airborne transmission of Mycobacterium tuberculosis: implications for control of tuberculosis in the HIV era. Clin Infect Dis. 2007;44(10):1358–60.

21. Escombe AR, Oeser C, Gilman RH, Navincopa M, Ticona E, Martinez C, et al. The detection of airborne transmission of tuberculosis from HIV-infected patients, using an in vivo air sampling model. Clin Infect Dis. 2007;44(10):1349–57.

22. Hatherill M, Tait D, McShane H. Clinical Testing of Tuberculosis Vaccine Candidates. Microbiol Spectr. 2016;4(5).

23. Bowling JD, O’Malley KJ, Klimstra WB, Hartman AL, Reed DS. A Vibrating Mesh Nebulizer as an Alternative to the Collison Three-Jet Nebulizer for Infectious Disease Aerobiology. Appl Environ Microbiol. 2019;85(17).

24. Zhang G, David A, Wiedmann TS. Performance of the vibrating membrane aerosol generation device: Aeroneb Micropump Nebulizer. J Aerosol Med. 2007;20(4):408–16.

25. Ault A, Zajac AM, Kong WP, Gorres JP, Royals M, Wei CJ, et al. Immunogenicity and clinical protection against equine influenza by DNA vaccination of ponies. Vaccine. 2012;30(26):3965–74.

26. Roy CJ, Reed DS. Infectious disease aerobiology: miasma incarnate. Front Cell Infect Microbiol. 2012;2:163.

27. Fears AC, Klimstra WB, Duprex P, Hartman A, Weaver SC, Plante KS, et al. Persistence of Severe Acute Respiratory Syndrome Coronavirus 2 in Aerosol Suspensions. Emerg Infect Dis. 2020;26(9).

